# CASi: A multi-timepoint scRNAseq data analysis framework

**DOI:** 10.1101/2023.08.16.553543

**Authors:** Yizhuo Wang, Christopher R Flowers, Michael Wang, Xuelin Huang, Ziyi Li

## Abstract

Single-cell RNA sequencing (scRNA-seq) technology has been widely used to study the differences in gene expression at the single cell level, providing insights into the research of cell development, differentiation, and functional heterogeneity. Various pipelines and workflows of scRNA-seq analysis have been developed but few considered multi-timepoint data specifically. In this study, we develop CASi, a comprehensive framework for analyzing multiple timepoints’ scRNA-seq data, which provides users with: (1) cross-timepoint cell annotation, (2) detection of potentially novel cell types emerged over time, (3) visualization of cell population evolution, and (4) identification of temporal differentially expressed genes (tDEGs). Through comprehensive simulation studies and applications to a real multi-timepoint single cell dataset, we demonstrate the robust and favorable performance of the proposal versus existing methods serving similar purposes.

## Background

In recent years, the emergence of the single-cell RNA sequencing (scRNA-seq) technique has enabled researchers to study cellular compositions and transcriptomic profiles with unprecedented precision. It has shown clinical utilities in many types of diseases, such as cancers [1, 2], Alzheimer’s disease [3, 4] and the SARS-CoV-2 infection [5, 6]. Specialized computational methods have been developed to identify differentially expressed genes across cell populations, find functionally distinct cell clusters, investigate the therapeutic effect of an external intervention, and more [7].

After finishing the preprocessing and quality control steps, the downstream analysis of scRNA-seq data includes dimension reduction, unsupervised clustering, differential expression analysis and so on [8]. The popular pipelines for these analyses include scran [9], Seurat [10], and SINCERA [11]. Recently, a number of supervised cell type annotation algorithms which can potentially identify novel cells have been developed, including scmap [12], CHETAH [13], and singleR and CAMLU [14]. In addition to recognizing the cell types, detecting differentially expressed genes (DEGs) is another common task in scRNA-seq analysis and lots of methods have been developed for this task. Simple statistical tests (i.e., the t-test and the Wilcoxon Rank Sum test) are used in pipelines such as Seurat. But more complex model-based frameworks have been shown to achieve better performance [15], e.g., DESeq2 [16] and MAST [17].

Although many existing methods have been developed, almost all of them focus on analyzing data collected from cross-sectional studies, i.e., the scRNA-seq experiments are performed on samples collected at a single time point. In the setting of studying disease development and progression, following patients for a period of time and collecting data from continuing experiments is actually a natural choice. Recent years have witnessed a growing number of multi-timepoint studies. For example, Ravindra et al. studied the SARS-CoV-2 infection by performing scRNA sequencing experiments on infected human bronchial epithelial cells at four time points to track patients’ immune responses to the virus [18]. In another study, Zhang et al. followed patients with ovarian cancer for five years and collected tissue samples before and after chemotherapy to study their stress-promoted chemoresistance [19]. Multi-timepoint scRNA-seq experiments provide a powerful tool for studying the dynamics of gene expression in individual cells and how it changes in response to various stimuli or disease conditions. At present, there are no available computational tools to comprehensively analyze multiple timepoints’ scRNA-seq data. Most of the current transcriptomic single-cell studies with the design of different timepoints still use methods and workflows, such as Seurat, which do not specifically take the time changes into consideration [18, 20]. Ramazzotti et al. proposed a framework called LACE that processes Single-cell mutational profiles, which are generated by calling variants from scRNA-seq data collected at different time points [21]. However, LACE specifically focuses on building somatic mutational profiles and reconstructing longitudinal clonal tree in tumor cells, not a general framework designed for multi-timepoint scRNA-seq data.

There are several reasons why cross-timepoint data can be more challenging to analyze compared with cross-sectional data. First, samples from the same subject are likely to be correlated. The information from previous time points are generally helpful for analyzing the data from later time points, and this should be considered for information borrowing. Such correlation also complicates differential signal detection. Second, new cell types may emerge over the experimental course. Ignoring the potential for new cell types can lead to inaccurate cell annotation and misleading results. Such newly emerged cells may also be the key to disease progression and treatment outcome, and thus they should be highlighted in the analysis procedures. Third, the single cell data are usually collected gradually over a long period of time. It is preferable for the analysis pipeline to take such collection schedules into consideration, when able, to allow for stepwise analysis and the incorporation of future data without changing the current results dramatically.

To address these challenges, we develop a comprehensive framework specifically to analyze scRNA-seq data from multi-timepoint experiments. Similar to other pipelines, the first step of our method is to annotate cell labels. But our method allows for information borrowing from earlier time points, as well as step-wise analysis. Our framework implements a neural network classifier to assign cell labels in a supervised way, and it can achieve high accuracy in the cross-timepoint setting. Nevertheless, one challenge faced by all the supervised annotation methods is identification of the novel cell type, which is defined as a cell type that is not present in the initial or earlier time points and that only exists in the newer collections. Along with the supervised annotation, we propose and implement a novel methodology pipeline in the framework to identify new cell types that have emerged over time. Once the annotation is done, we provide visualizations to illustrate the cell population evolution. It should be Additionally, we add one key downstream analysis, temporal differential expressed gene (tDEG) detection, to our framework. The aim is to identify genes with wildly increasing/decreasing behavior over time and with different changing patterns from group to group. For example, our framework is able to select the genes whose expression increases over time for responders but decreases over time for non-responders. The whole framework is named the Cross-timepoint Analysis of Single-cell RNA sequencing data (CASi) method.

## Results

### An overview of CASi

CASi takes scRNA-seq data collected from different timepoints as the input (Fig. 1). To simplify the discussion, we consider an scRNA-seq dataset collected at three time points: *t*_0_, *t*_1_, and *t*_2_. Assuming *t*_0_ data is pre-labeled, i.e., *t*_0_ data is provided with cell types, the labels can be obtained using existing unsupervised or supervised approaches for cross-sectional data. The first step of CASi is to perform a supervised annotation for unlabeled data *t*_1_ and *t*_2_ using an artificial neural network classifier. However, if the labels for the *t*_0_ data are not provided, CASi instead will perform an unsupervised clustering and use the clustering number as the cell type labels for *t*_0_ data. Then the classifier can be trained with *t*_0_ data and applied to annotate *t*_1_ and *t*_2_ data. Here we prefer an artificial neural network over other machine learning methods because a few recent works [22, 23] showed its superior accuracy and favorable computational performance in the task of cell type annotation.

**Figure 1.**
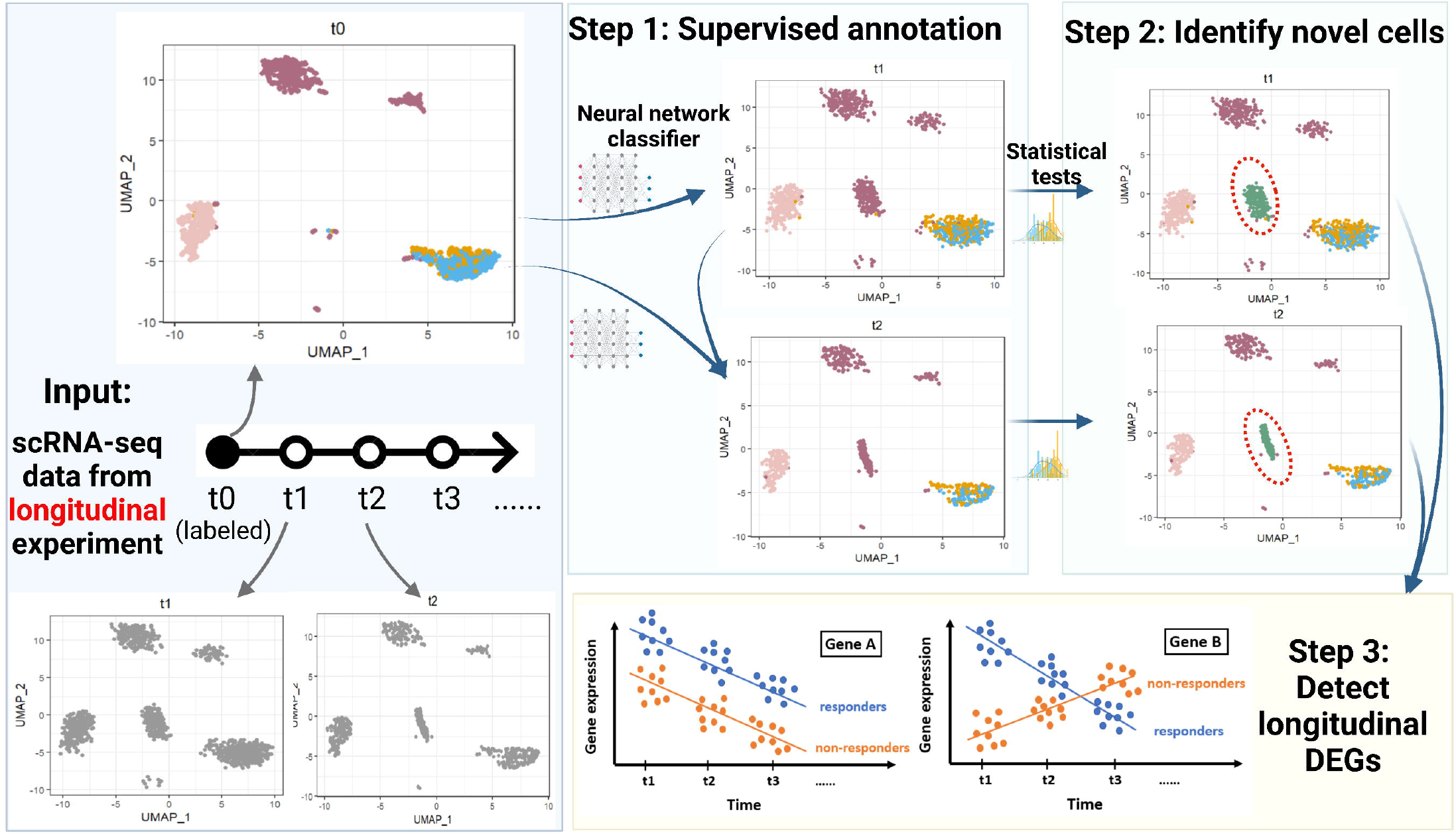
An overview of the CASi framework. The input is scRNA-seq data from different timepoints’ experiments. CASi mainly consists of three steps: 1) cross-time points cell annotation, 2) detection of potential novel cell types, 3) identification of temporal differentially expressed genes

The second step of CASi is to identify any novel cell types that have emerged over time. When data of interest is collected from tumor samples, it is highly possible that the tumor cells will differentiate and new cell types (e.g., a distinct subclone of tumor cells) might appear. Assuming the cells of *t*_1_ and *t*_2_ data are a mixture of known and unknown cell types, we have designed a computational pipeline to distinguish these new cell types from existing cell types. The pipeline starts with a feature selection procedure to select a smaller set of informative features, followed with a dimension reduction procedure to better extract information from these features. Then we use the two-sample t-test to compare the correlations between features in known cell types and features in unknown cell types, and thus we are able to identify novel cells. Intuitively, after using the neural network to assign cell types for *t*_1_ and *t*_2_ data, if the cells actually belong to a new, unknown cell type, the levels of similarity with the known cell types will be lower than all of the other cells. Additionally, it should be noted that, CASi allows for the detection of multiple novel cell types. We achieve this by providing users with both the cluster-labeled UMAP (Uniform Manifold Approximation and Projection) plot and the correlation-labeled UMAP plot, which visually, directly reveals the possibility of multiple novel cell types.

The forth and final step of CASi is to perform differential analysis tailored to multi-timepoint scRNA-seq data. We combine a generalized linear model with iterative feature selection to select genes that have apparent increasing/decreasing behavior over time and genes that behave differently along time in different groups. More details of CASi are presented in the Methods section.

### Simulation settings

The simulation data are generated by assuming three sampling time points: *t*_0_, *t*_1_, *t*_2_. To fully evaluate our method, we designed three scenarios: 1) *t*_0_, *t*_1_, and *t*_2_ data contain the same cell types but with different cell type compositions; 2) a cell type in *t*_0_ disappears in *t*_1_ and *t*_2_ data, i.e., *t*_0_ data have one more cell type than *t*_1_ and *t*_2_ data; 3) a new cell type appears in *t*_1_ and *t*_2_ data, i.e., *t*_1_ and *t*_2_ data have one more cell type than *t*_0_ data. We obtain a publicly available dataset of peripheral blood mononuclear cells (PBMC) [24], containing more than 60,000 sorted cells from eight immune cell types. We randomly extract cells from five cell types and use different cell type compositions for different scenarios. Detailed settings of the three scenarios can be found in the Method section. In this study, we compare our framework to the existing methods, including supervised clustering methods such as scmap [12], CHETAH [13], and scPred [25] to evaluate the annotation accuracy. We also compare the proposal with other differential expression analysis methods, such as DESeq2 [16], MAST [17], and the Wilcoxon Rank Sum test offered in Seurat [26], to evaluate the performance of tDEG detection.

### CASi facilitates cross time points cell annotation with high accuracy

CASi is suitable when data of interest are collected from multiple timepoints as it borrows information across time points and allows for accurate cell annotation when data becomes gradually available. Using the neural network classifier, CASi achieves a high accuracy when mapping the cell labels of the initial time point onto later time points. The accuracy is defined as (number of correctly labeled cells at time *t*_1_ and *t*_2_) / (total cell number of *t*_1_ and *t*_2_ data). In addition to accuracy, we use adjusted rand index (ARI) to assess the performance of the neural network classifier, which is a widely used metric to evaluate clustering performance. Rand Index looks at similarities between any two clustering methods and ARI is the corrected-for-chance version of the Rand index [27]. The accuracy and ARI of each scenario based on 200 Monte Carlo experiments are shown in Fig. 2.

**Figure 2.**
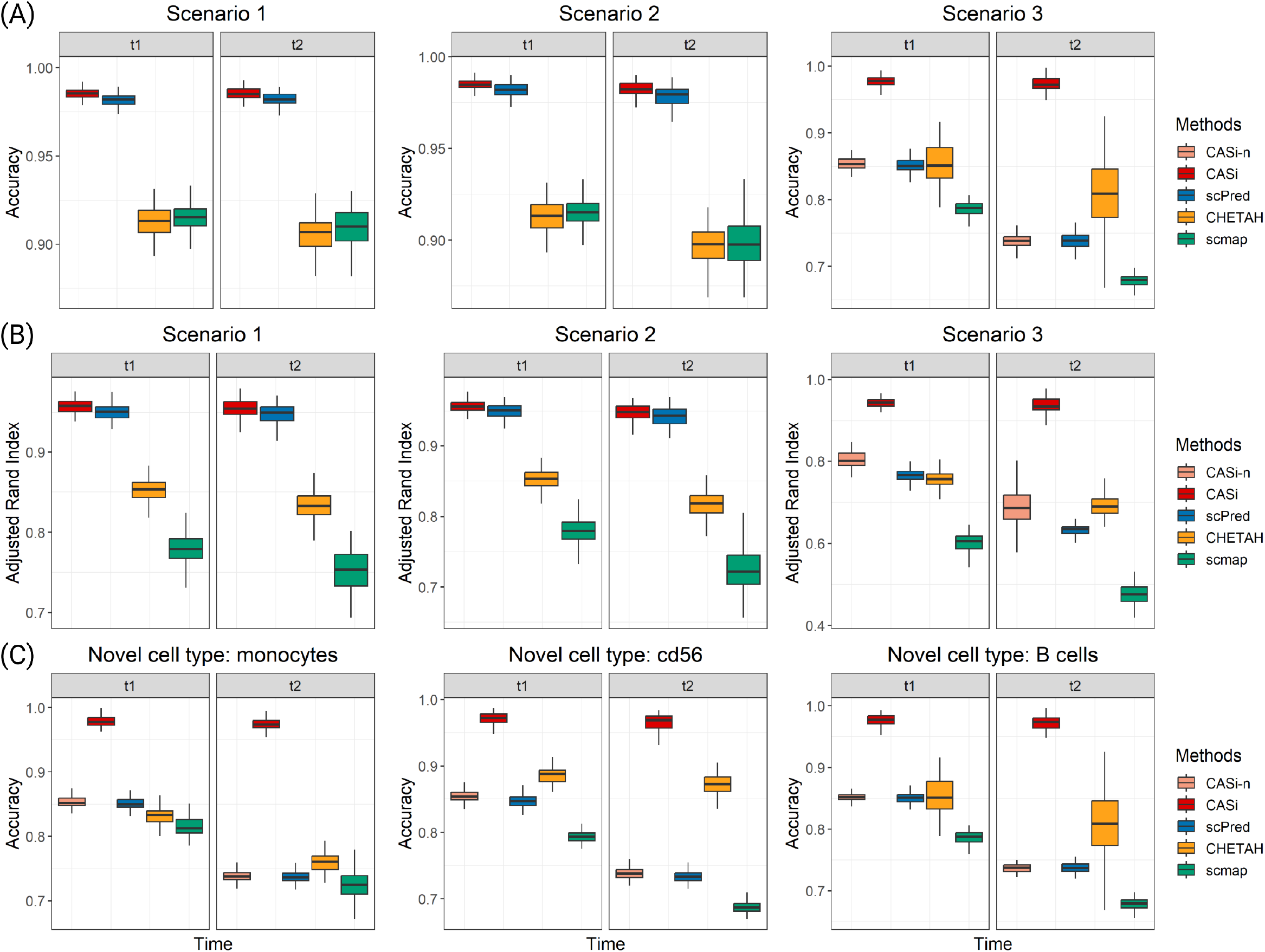
Results of PBMC simulation study based on 200 repetitions. Scenario 1: *t*_1_ and *t*_2_ data have different cell populations compared with *t*_0_ data; scenario 2: one cell type disappears in *t*_1_ and *t*_2_ data; scenario 3: one new cell type appears in *t*_1_ and *t*_2_ data. Note that for scenario 3, an additional method is included. CASi-naive (CASi-n) refers to only the neural network classifier without our follow-up pipeline. Panel A: accuracy of the cross-time points cell annotation. Panel B: ARI of the cross-time points cell annotation. Panel C: accuracy of scenario 3 using different cell types as the new cell types emerged in *t*_1_ and *t*_2_.

It can be observed that, for scenario 1 and 2, our method has the highest accuracy and the highest ARI, but the advantage is not significant when compared to the scPred method. For scenario 3, we report the performance of CASi-n, which refers to only the neural network classifier, and the performance of CASi, which refers to the neural network classifier in alignment with the identify-novel-cell pipeline. As we expected, the disappearance of one cell type (scenario 2) does not affect the classifier’s performance, while the appearance of one novel cell type (scenario 3) causes trouble to the classifier. After our pipeline identifies and labels the novel cells, both the accuracy and the ARI improve significantly, and CASi demonstrates state-of-art performance when novel cell types appear. For scenario 3, we vary the setting of scenario 3 and use different cell types as the novel cell type. In Panel C, we report the accuracy of these settings and observe a similar pattern that CASi outperforms other existing methods.

### CASi addresses possibility of novel cell types appeared in later time points

CASi uses the levels of similarity between the known cell types and new cells to identify potential novel cell types that may have appeared at later time points. The full pipeline of identifying novel cells is described in the Method section. We provide users with the Uniform Manifold Approximation and Projection (UMAP) plots displaying cell type clusters and the UMAP plots displaying the correlation. Fig. 3 Panel A and B shows the UMAP plots of *t*_1_ and *t*_2_ data, respectively. The purple group represents novel cells that appeared in *t*_1_ and *t*_2_. And we can see that the correlation between this new cell type and the existed cell type is very low. For scenario 3 in the simulation, we only assume one new cell type appears. When *> one* new cell type appears in later time points, users will be able to recognize them separately as new cells will be clustered into multiple groups on the UMAP plots.

**Figure 3.**
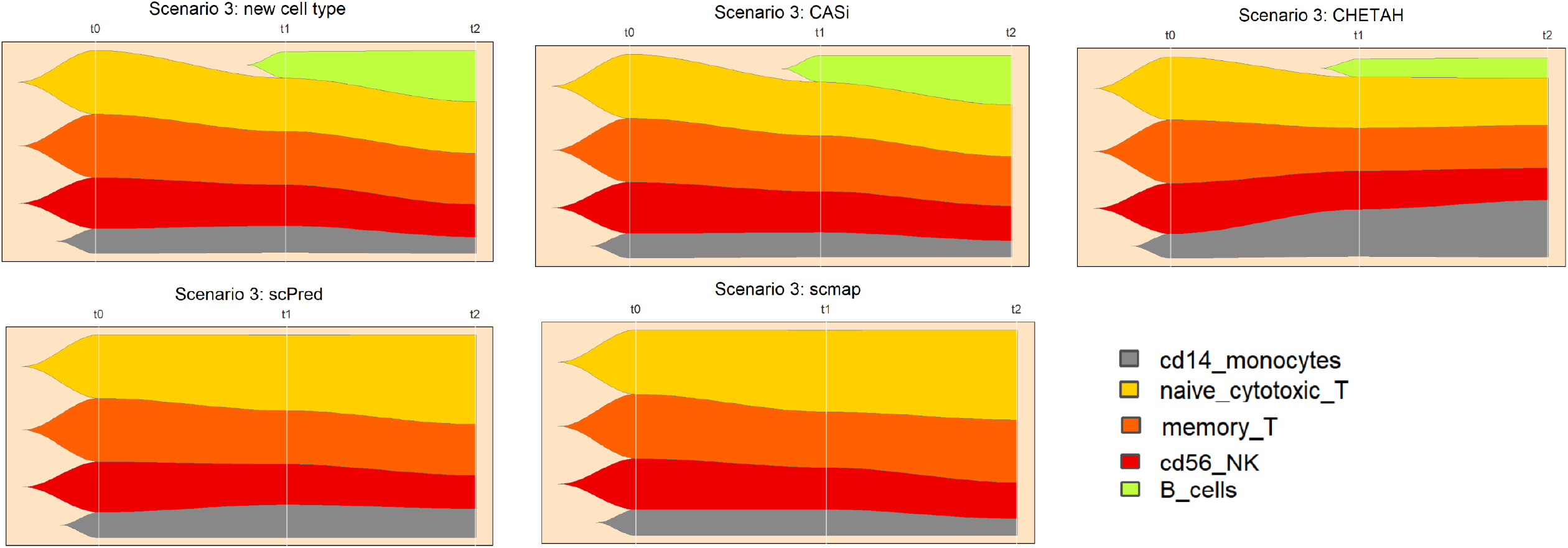
UMAP plots of the PBMC simulation study: intuition behind identifying novel cell types. Panels A and B show *t*_1_ and *t*_2_ data, respectively. In both panels, the left figure is the clustering of true cell labels, the middle figure is the clustering that highlights the novel cell type group, and the right figure is the clustering of correlation levels between existing cell types in *t*_0_ data and new cell types in *t*_1_/*t*_2_ data

### CASi provides visualization of temporary cell type evolution

To track the changes of the complex cell population, an appropriate form of visualization is of importance. We use a combination of UMAP and fish plots to illustrate the dynamic changes in the cell type proportions over time. Traditional UMAP/scatter plots visually show how separable the clusters/cell types are using selected features. The fish plot developed by Miller et al. is a relatively new tool displaying changes in populations of cells over time [28], which is a great tool to visualize the temporal cellular proportion evolution. In fish plots, each color represents one cell type; the first plot in Fig. 4 shows the true cell type population of Scenario 3, where a novel cell type appears in *t*_1_ and *t*_2_. We can see that the CASi prediction is highly similar to the ground truth and able to identify most novel cells, while other methods either only identify a small portion of novel cells or fail to capture any novel cells.

**Figure 4.**
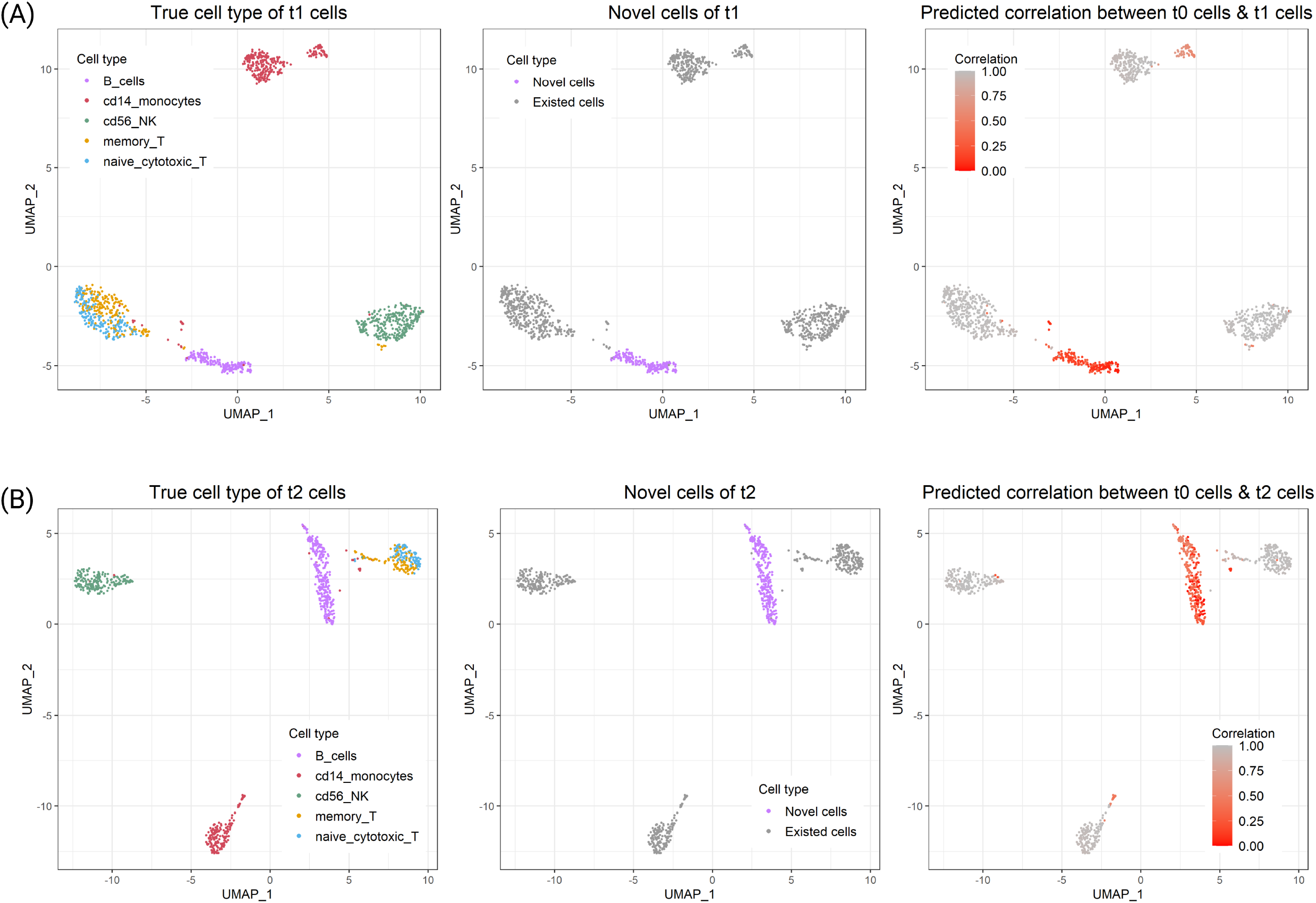
Fish plots of scenario 3 in which a novel cell type appears. We use B cells as the novel cell type which shows in green. The first figure is the ground truth of cell population and CASi (the second figure) is able to annotate the cells with high accuracy compared with other methods

### CASi identifies tDEGs

A natural interest in multi-timepoint scRNA-seq data analysis is to detect genes that change vividly over time, and different groups might have different changing directions. We refer to this kind of gene as tDEGs. To simulate tDEGs, we need to add both group effects and time effects. We design three levels of time effect: weak, medium, and strong. And since differential analysis is generally performed per cell type, we only use monocytes in this simulation. We randomly select 2000 genes and 900 monocyte cells from the PBMC dataset. Among 2000 genes, 300 genes are randomly selected to be tDEGs. Detailed settings are described in the Method section.

We use true discovery rate (TDR), which is a measure of accuracy when multiple hypotheses are being tested at once, to evaluate this step. We compare the performance of CASi and existing methods including DESeq2 [16], MAST [17], and the Wilcoxon Rank Sum test offered by the FindMarker() function in Seurat [26]. Fig. 5 shows that for all three time effect levels, CASi has the highest TDR at different thresholds, which means that CASi is able to capture tDEGs at the greatest extent. When the time effect increases, the TDR of CASi increases as well, but the TDR of other methods decreases.

**Figure 5.**
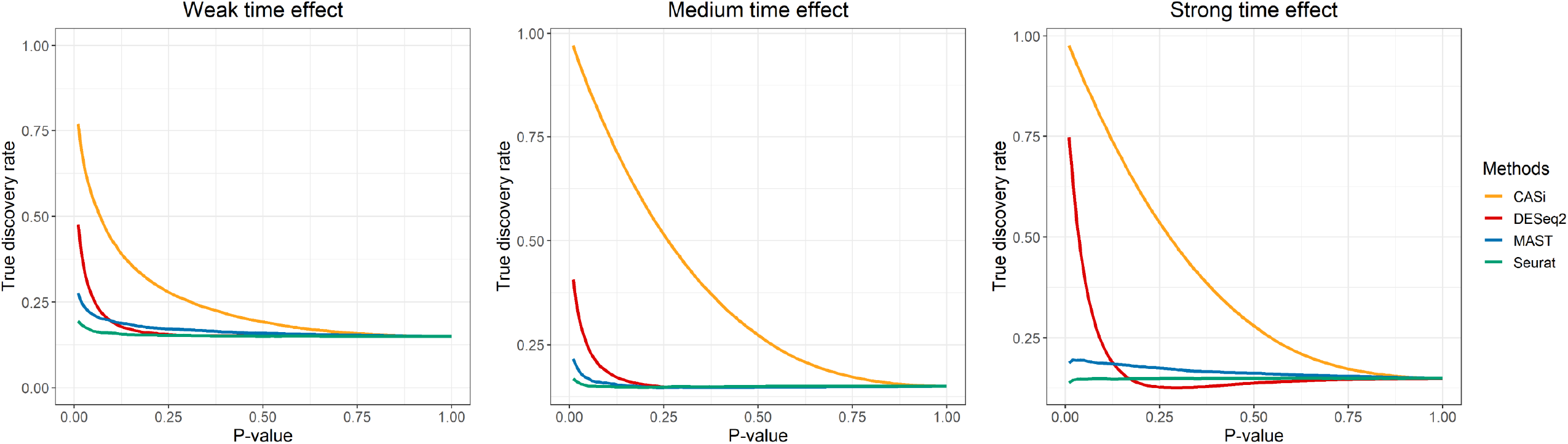
True discovery rate of identifying temporal differentially expressed genes. The results are averaged based on 200 repetitions. Three simulation settings are being considered here: weak time effect (left), medium time effect (middle), strong time effect (right)

### Apply CASi to a real-world multi-timepoint scRNA-seq data

We apply CASi to a real-world mantle cell lymphoma (MCL) dataset [29] where all patients received ibrutinib, the current standard of care treatment for MCL, but responded differently to the treatment. Three patients are ibrutinib-responsive (patients V, C, and D) and two patients are non-responsive (patients B and E). The MCL dataset requires cross-timepoint analysis: it includes measurements of 21 specimens collected at baseline, during treatment, and/or at disease remission/progression. Since the number of measurements and the timing of measurements vary from patient to patient, we manually binarize the time variable into two groups: pre-treatment and post-treatment, which also aligns with the analysis in the original paper of the MCL data.

#### Cross time points annotation

We again use accuracy and ARI to compare CASi with existing methods. From Fig. 6, Panel B, we can see that when mapping the cell labels of pre-treatment data to the cell labels of post-treatment data, CASi achieves the highest accuracy and the highest ARI. Panel A displays the true cell composition of post-treatment and the predicted cell composition by CASi, the two of which are visually identical. In Panel C, we show UMAP plots of cell types (left) and correlation (right). The tumor cells are separated into two clusters and the correlation between tumor cells in posttreatment data and tumor cells in pre-treatment data are not the same for these two clusters. The cluster with a correlation of 0.5-1 is more similar to the tumor cells in pre-treatment data, while the cluster with a correlation of 0-0.25 is very different from the tumor cells in pre-treatment data. This is a very interesting observation since tumor cells appear to be more genetically unstable than normal cells and may evolve over the treatment course [30]. Over time, tumor cells divide more rapidly and become less dependent on signals from other cells. This is probably why we observe two clusters for post-treatment tumor cells with two levels of correlation.

**Figure 6.**
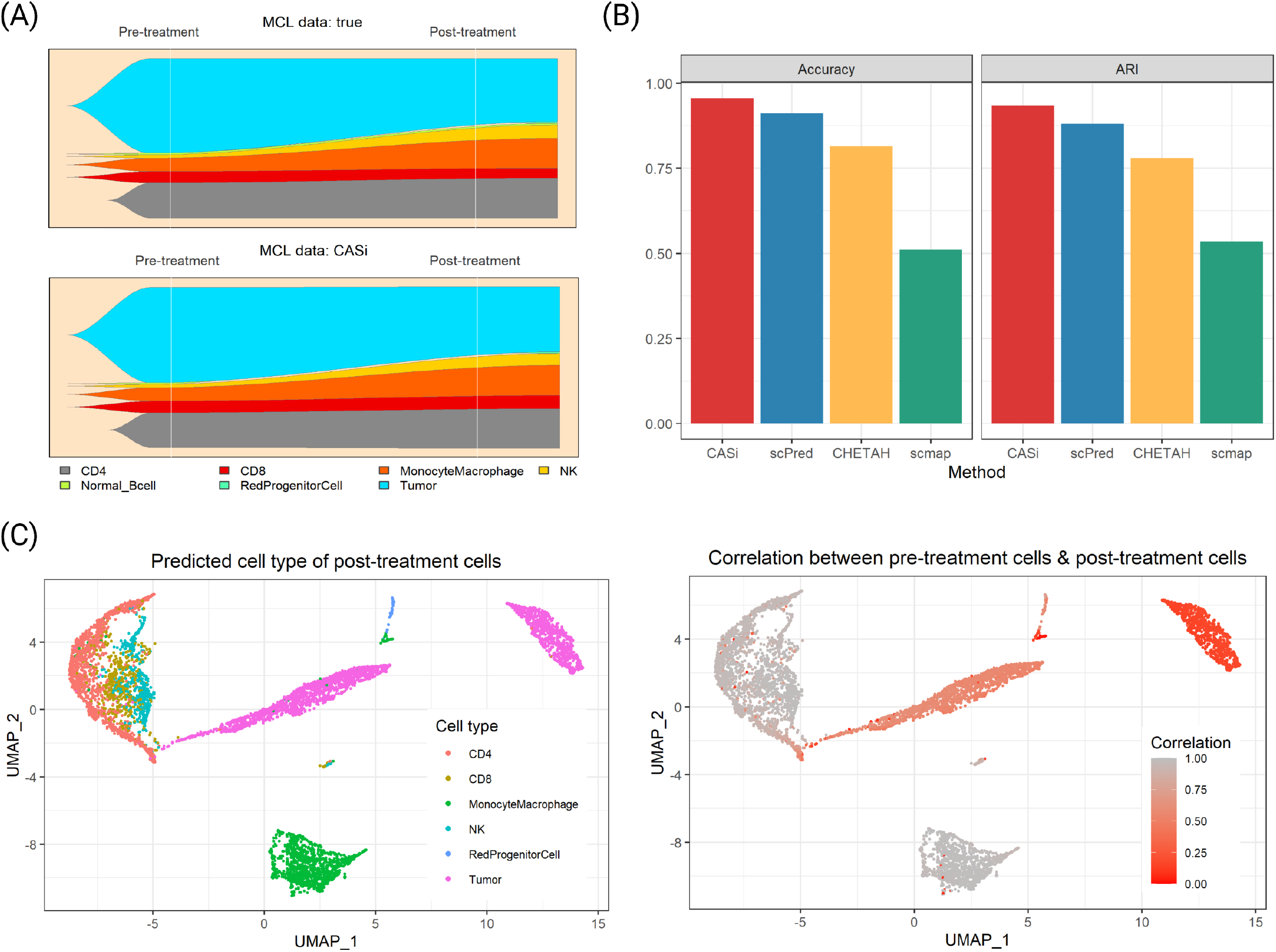
Annotation results of real-world data application. Panel A shows the fish plots of the true cell population (top) and the annotated cell population of CASi (bottom). Panel B shows the accuracy (left) and adjusted rand index (right) of the supervised clustering step. Panel C shows the clustering of cell labels (left) and the clustering of correlations between pre-treatment cells and post-treatment cells (right) in which red represents a low correlation level

#### Identification of tDEGs

tDEGs refer to the genes that change vividly over time, and the changing patterns are distinct for different groups. We apply CASi and Seurat at the same time to this dataset to find tDEGs. Using p = 0.05 as a cutoff, among a total of 1469 genes, CASi identifies 268 tDEGs and Seurat identifies 75 DEGs. There are 21 overlapped genes for tDEGs found by CASi and DEGs found by Seurat. CASi is able to identify a few tDEGs that Seurat failed to capture their significance. In Fig. 7 Panel A, we illustrate the two top tDEGS identified by CASi but not Seurat, IFITM2 (the top plot) and IFITM3 (the bottom plot), and we draw their gene expression patterns for all patients. Patients B and E are non-responders and are drawn in blue; patients C, D, and V are responders and are drawn in red. Clearly, for responders, the gene expression of the identified genes increases after the ibrutinib treatment, while for non-responders, the gene expression decreases after the ibrutinib treatment. This finding is supported by a recent study in which Lee et al. discovered that deletion of IFITM3 in MCL cells reduced competitive fitness and proliferation, and the upregulation of IFITM2 might suggest a compensation mechanism for the loss of IFITM3 [31].

**Figure 7.**
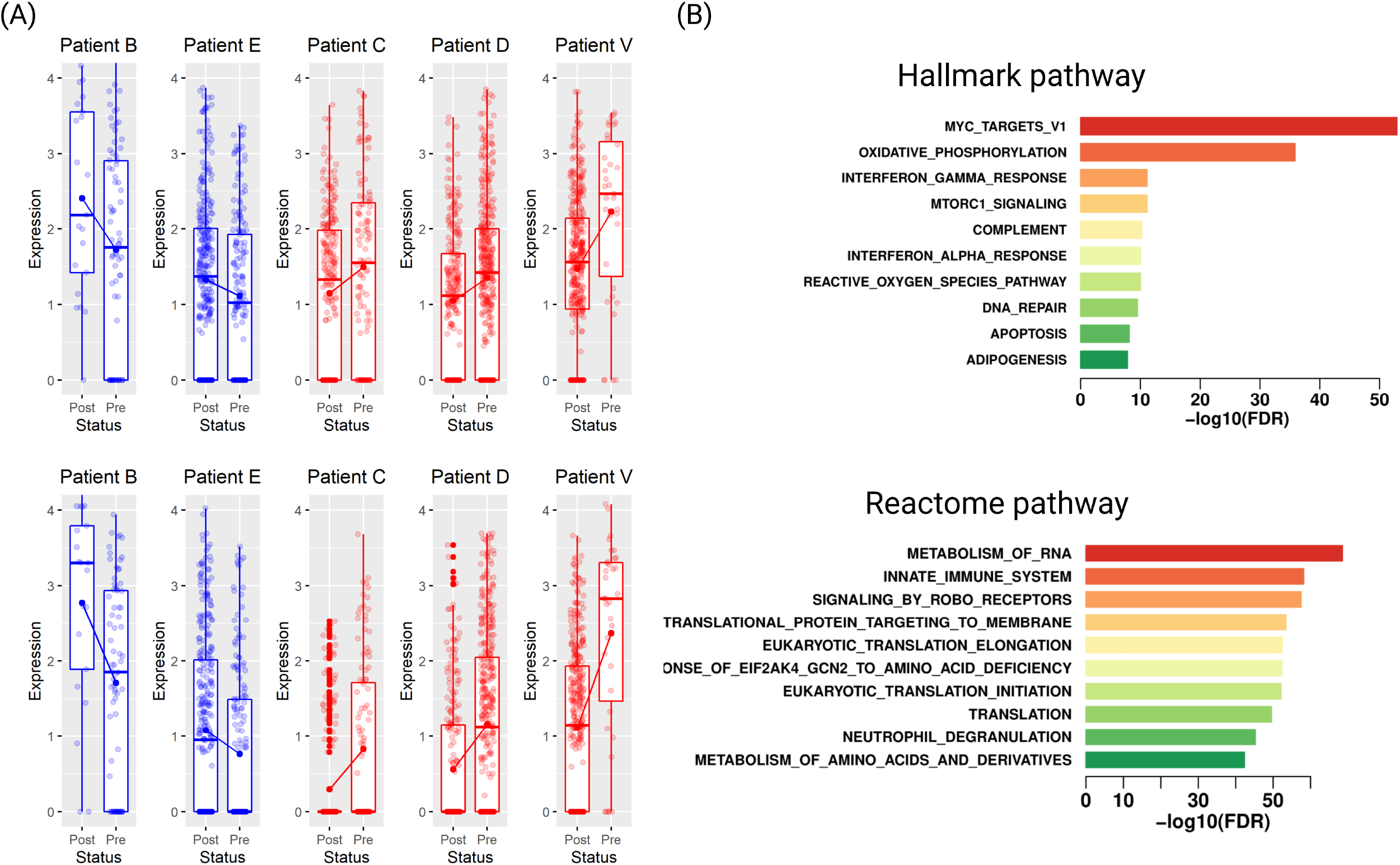
Differential expression analysis results of real-world data application. Panel A shows the changing gene expression profiles of IFITM2 (top) and IFITM3 (bottom), which are temporal differentially expressed genes identified by CASi but not Seurat. Non-responders, including patients B and E, are shown in blue; responders, including patients C, D, and V, are shown in red. Totally opposite patterns can be seen for responders and non-responders. Panel B shows the results of enrichment analysis using 500 temporal differentially expressed genes identified by CASi: hallmark pathways (top) and reactome pathways (bottom)

Additionally, using the top significant genes selected by CASi, we perform enrichment analysis, which identifies biological pathways that are enriched in this gene list more than just by chance. Fig. 7 Panel B shows the Hallmark pathway results (top) and the Reactome pathway results (bottom). For the top two hallmark pathways, MYC and oxidative phosphorylation, multiple studies have reported that MYC is frequently expressed in MCL, and targeting MYC provide a novel therapeutic strategy for MCL patients [32, 33]; multiple studies have found that the MCL cancer cells can be effectively targeted with a small-molecule inhibitor of oxidative phosphorylation as a therapeutic strategy [34, 35].

For other hallmark pathways, there also are studies that have reported their close relationships with MCL. For example, some researchers have reviewed the abnormality of interferon gamma inducible protein 16 (IFI16) expression in MCL and the involvement of Complement related to the drug-resistance mechanism of MCL [36, 37]. For reactome pathways, several researchers have findings that match our own in that RNA metabolism pathway is significantly enriched in MCL [38] and that the ibrutinib resistance of MCL relates to the receptors initiating the innate immune system [39, 40]. The drug, lenalidomide, which has demonstrated strong therapeutic effect when treating MCL, is a immunomodulatory drug with effects on the innate immune system [41]. Additionally, multiple studies have reported that the expression of eukaryotic translation initiation factors may play an important role in the development of MCL, and that they are highly correlated with their tumor grade [42, 43, 44].

## Discussion

Although a series of clustering and annotating methods have been developed for scRNA-seq data, these methods mostly focus on cross-sectional studies. A systematic analysis tool designed specifically for cross-timepoint scRNA-seq datasets is lacking. With a rapidly growing number of studies conducting experiments at different timepoints, there is a great need for methods to analyze multi-timepoint single-cell data. In this study, we present CASi, the first framework to provide a full analysis pipeline for analyzing scRNA-seq data from multi-timepoint designs, ultimately creating an informative profile of dynamic cellular changes.

It should be noted that the proposed method has a fundamentally different goal from existing pseudo-time trajectory inference methods, such as Monocle [45], SCUBA [46], and TSCAN [47]. Although both CASi and pseudo-time inference methods can take experimental time into consideration, pseudo-time methods focus on the innate cell differentiation/cell-state transition process, while CASi provides a more general view of the dynamic changes of the cell population over multiple timepoints. Moreover, regardless of the term “pseudo-time ordering,” the ordering of cells by those methods can be any type ordering and might not have a time interpretation, such as with spatial order [48].

The first step of CASi uses the neural network classifier to achieve cross-time points cell annotation with high accuracy. And as a supervised learning method, it efficiently avoids the overclustering issue. Overclustering often appears in unsupervised clustering. When the total number of cells increases, the cells of one type will be separated into two or even more clusters. Using the same scenario settings of the simulation, we compare ARI of unsupervised clustering implemented in the Seurat package with ARI of CASi using supervised clustering. The results of three scenarios are shown in the supplementary file. It can be observed that our method’s ARI increases with the cell number increasing, while the Seurat ARI decreases with the cell number increasing. This indicates that the supervised clustering method of CASi avoids the overclustering problem.

However, supervised clustering methods also have a universal disadvantage when dealing with novel cell types. When the new/unknown cell types appear only in the testing data, the classifier will not be able to distinguish them and will assign these cells to an existing but wrong cell type. In continuing experiments, with the progression of the tumor, some existing cell types might disappear and some new/unknown cell types that are not present at the beginning might appear later. Another innovation of our work is the detection of novel/unknown cell types that emerge over time. We designed a pipeline using correlation and t-test in a robust way to distinguish novel cell types. We have demonstrated state-of-the-art power in both simulation studies and real-world data application compared with existing methods serving a similar purpose.

Differential expression analysis is one of the most common tasks in scRNA-seq studies, and it is also a critical step in CASi. Several methods have been proposed to detect DEGs, including BPSC [49], MAST [17], and Monocle [50]. These methods are designed for two group comparisons using scRNA-seq data, and they may have difficulty accounting for the time effect and interaction terms across different time-points’ scRNA-seq data. CASi allows users to detect tDEGs, which refers to genes that express wildly changing behavior over time. To find tDEGs, we combine the generalized linear model with iterative feature selection. For each gene, a p-value is obtained from the GLM model to indicate evidence of differential expression. We initially started the analysis with the generalized linear mixed model (GLMM) to account for subject’s random effect. However, GLMM is computationally expensive, and looping through each gene is infeasible in most scenarios. We then turned to GLM and have found that the regression analysis results, i.e., the coefficients and p-value, are very similar between GLM and GLMM models in the explored settings. Thus, taking into consideration the model complexity and the computational cost, we eventually chose GLM to model the multi-timepoint gene expression count data. Current methods can be extended in a few ways. First, the novel cell detection framework may not work well when the novel cell is transcriptomically similar to the known cell types. In this scenario, it may be beneficial to incorporate other information, such as copy number variation or mutation data, into consideration. Second, the current cell type annotation step pools the data from all the samples together for training and applying the neural network. It is possible that the cell populations are quite distinct for subjects from different conditions. As a result, it may be beneficial to perform the annotation in subjects within the same condition if sample size allows. Finally, the current GLM framework can only detect linear effect, i.e., the linear change of gene expression by time or the interaction of time and condition. It helps if the model can be extended to allow the detection of nonlinear changes.

## Conclusions

With the growing number of scRNA-seq studies collecting data in a continuous manner, there is a lack of bioinformatics tools specifically designed to take time effect into consideration. We propose CASi, the first comprehensive pipeline for analyzing and visualizing multi-timepoint scRNA-seq data. CASi is advanced in mapping cell labels across multiple time points, identifying novel cell types, visualizing cell population evolution, and detecting temporal differentially expressed genes. We hope our framework will fill in the gap and facilitate prospective single cell studies.

## Methods

### Artificial neural networks (ANNs)

Denote the scRNA-seq expression matrix of the training data by ***X***_0_ where ***X***_0_ is a *p* by *n*_0_ matrix with *p* being the total number of measured genes and *n*_0_ is the number of cells, and the corresponding training cell label, ***Y*** _0_, is a *n*_0_ by 1 vector. Similarly, denote the expression matrix of the testing data by ***X***_1_, which has dimensions *p* by *n*_1_ and the labels by ***Y*** _1_. After the standard min-max normalization, we select the top 2000 most variable genes from *X*_0_ and the same set of features from ***X***_1_. Next, using Keras [51], an open-source software library that provides a Python interface for artificial neural networks, we train a neural network model with one input layer, one output layer, and three hidden layers:

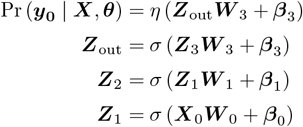

The parameter set *θ* = {***W*** _0_, ***W*** _1_, *…*, ***W*** _3_, ***β***_0_, ***β***_1_, *…*, ***β***_3_, ***β***_out_, ***W*** _out_ } will be estimated during the training process. And *z*_*l*_ for *l* = {1, 2, 3} are the hidden neurons with corresponding weight ***W*** _1_, and bias ***β***_1_.*σ*(*·*) is the activation function, which can be a sigmoid, a rectified linear unit (ReLU), a hyperbolic tangent, etc. We choose to use the ReLU function in our hidden layers because the neural networks based on an ReLU function are generally easier to train and can avoid the vanishing gradient problem during optimization [52]; it is mathematically expressed as *σ*_ReLU_(*x*) = max(*x*, 0). The SoftMax function will be used in the output layer. This is because the number of output categories is more than two, and it converts the values of the output layer into the predicted probabilities of each label. The number of neurons in the three hidden layers is selected as {256, 128, 64}. The model is trained using a stochastic gradient descent (SGD)-based algorithm with the mean squared error loss function **ℒ** (Pr(***y***_**0**_), *Y*_0_) = ∥Pr(***y***_**0**_) − *Y*_0_∥^2^. We use Adam as the optimization algorithm [53], and the mini-batch training strategy [54], which randomly trains a small proportion of samples and validates the rest of the samples in each iteration to improve training efficiency. By monitoring the loss, we implement the early stopping rule in Keras to avoid overfitting. Once the model performance stops improving for a couple epochs, the training process will stop. Additionally, to further prevent the overfitting issue, we add a dropout step with the dropping rate of 0.4 for each hidden layer to randomly drop units from the neural network during training [55].

### Identify novel cell types

Let *Y* ^*K*^, *K* = {1, …, *k* − 1, *k}* be the cell labels, and *Y* ^*k*^ is the novel cell type that only appears in the testing data *X*_1_. Then when annotating the cells in *X*_1_, all *Y*^*k*^ cells will be wrongly labeled as *Y* ^*K*^, *K* = {1, …, *k* − 1} by the neural network classifier. To address this issue, we design a pipeline. The key idea of our pipeline is that the similarity (correlation) of the new/unknown cell type with the known cell types will be different (usually smaller), which can be captured by the two-sample t-test. The final cell identities of the identified new cell types will be confirmed with external biological knowledge and expert opinions. The pipeline consists of the following steps:

1. Reduce dimension:

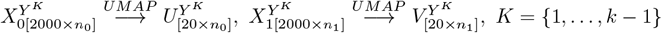
2. Obtain the mean of UMAP vectors for each cell type:

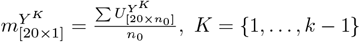
3. Obtain the Pearson correlation between cells in *X*_1_ and the mean:

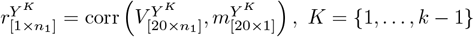
4. Re-cluster each cell type in *X*_1_ into two groups using the Louvain algorithm [56]:

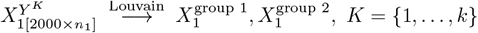
5. Apply two-sample *t* test to the two groups of cells using their cell type-specific correlation values. If significant, we will designate the group of cells that has a smaller mean correlation as the potential new cell type.
6. For each cell type *Y* ^*K*^, the t-test will assign a small group of cells as the new cell type. And in the end, we combine those significant groups of cells and annotate them as the new cell type *Y*_*k*_.

### Three scenarios of the simulation study for cross cell type annotation

Suppose we have scRNA-seq data collected at three different time points: *t*_0_, *t*_1_, *t*_2_. We design three simulation scenarios: 1) *t*_0_, *t*_1_, and *t*_2_ data contain the same cell types but with different percentages; 2) one cell type disappears in *t*_1_ and *t*_2_ data; one new cell type appears in *t*_1_ and *t*_2_ data. There are eight cell types included in the original dataset: CD4, B cells, CD14 monocytes, CD56 NK cells, memory T, naive T, naive cytotoxic T, and regulatory T. We randomly pick five of them to use for simulation studies: B cells, CD14 monocytes, CD56 NK cells, memory T, and naive cytotoxic T. We draw cells from each cell type without replacement and a total of 4500/4400/4300 cells are drawn for scenarios 1/2/3. Detailed cell type compositions of scenario 1, 2, and 3 are shown in Table 1.

**Table 1.**
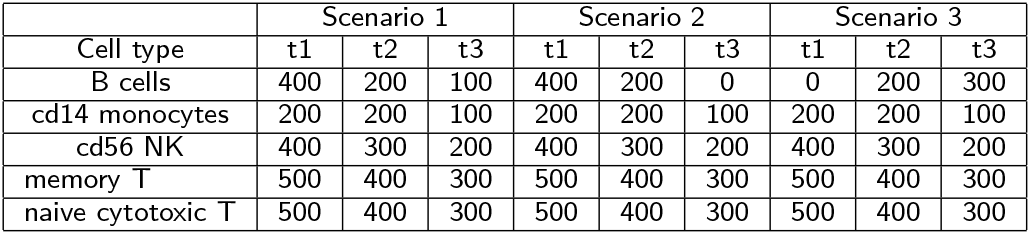
Simulation: cell type compositions.

### Simulation studies to evaluate the detection of temporal differential analysis

To evaluate the performance of identifying tDEGs, we generate multi-timepoint scRNA-seq data with a parametric model. For each gene, we build a negative binomial GLM model using a log link function. The negative binomial model is a generalization of the Poisson model such that the count *Y*_*i*_ still adopts a Poisson, but the expected count 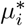 is a gamma-distributed random variable with mean *μ*_*i*_ and constant scale parameter *ω*, i.e., the mean and variance are not equal anymore. Mathematically, the count *Y*_*i*_ follows a negative-binomial distribution [57]:

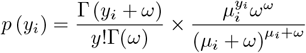

where the expected value is *E* (*Y*_*i*_) = *μ*_*i*_ and the variance is 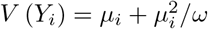. The covariates we use are the time variable, the strata of interest (e.g., the response status), and the interaction term of time and strata as independent variables, and the gene expression count is *Y*_*i*_. Note that for the existing methods, including DESeq2 [16], MAST [17], and the Wilcoxon Rank Sum test offered by the FindMarker() function in Seurat, only one binary covariate (i.e., the strata of interest) can be used in the analysis. This means that multiple covariates and the interaction term are not under consideration when finding tDEGs. And when we fit the negative binomial GLM model, both the regression coefficients and *ω* will be estimated by the method of maximum likelihood. For each model, the p-value of each term is obtained from testing the null hypothesis of its coefficient being equal to zero. Then for all genes, we extract the interaction terms’ p-values and use the Bonferroni correction to account for multiple-test effect. To evaluate this step, we conduct extensive simulations by manually creating tDEGs and calculating the true discovery rate. We extract cells from a publicly available PBMC dataset. Considering that different types of cells will have very different gene expression profiles, we only extract monocyte cells from the PBMC data to do simulation. We draw 2000 genes and 900 monocyte cells. Among 2000 genes, 300 genes are randomly chosen to be tDEGs.

First, we add group effect. Among 900 cells, half of the cells are assigned to be responders by multiplying the baseline gene expression by *Unif* (1.5, 2), i.e., a uniform distribution of min = 1.5 and max = 2, while the other half of the cells are assigned to be non-responders and their baseline gene expressions stay the same.

Next, we divided the whole data into three time points (*t*_0_, *t*_1_, and *t*_2_); namely we will have 150 responder cells and 150 non-responder cells at each time point. We manually assign 300 genes to be the tDEGs by adding the time effect. We design three levels of time effect: weak, medium, and strong. For the weak time effect scenario, the baseline gene expression of *t*_0_, *t*_1_, and *t*_2_ cells are multiplied by *Unif* (0.8, 1), *Unif* (1, 1.2), and *Unif* (1.2, 1.4), respectively. For the medium time effect scenario, the baseline gene expression of *t*_0_, *t*_1_, and *t*_2_ cells are multiplied by *Unif* (0.4, 0.6), *Unif* (1, 1.2), and *Unif* (1.6, 1.8), respectively. For the strong time effect scenario, the baseline gene expression of *t*_0_, *t*_1_, and *t*_2_ cells are multiplied by *Unif* (0.4, 0.6), *Unif* (1.2, 1.4), and *Unif* (2, 2.2), respectively.

## Appendix

### Availability of data and materials

The PBMC Single-cell expression data used for simulation study was obtained from the 10X website (https://support.10xgenomics.com/) [24]. The MCL Single-cell expression data used for real-world data application was downloaded from the European Genome-Phenome Archive (EGA) database with the accession code EGAS00001005019 [29]. Source codes for running CASi and installing the R package are available on GitHub (https://github.com/yizhuo-wang/CASi).

## Additional Files

### Additional file 1

**Supplementary figure:**
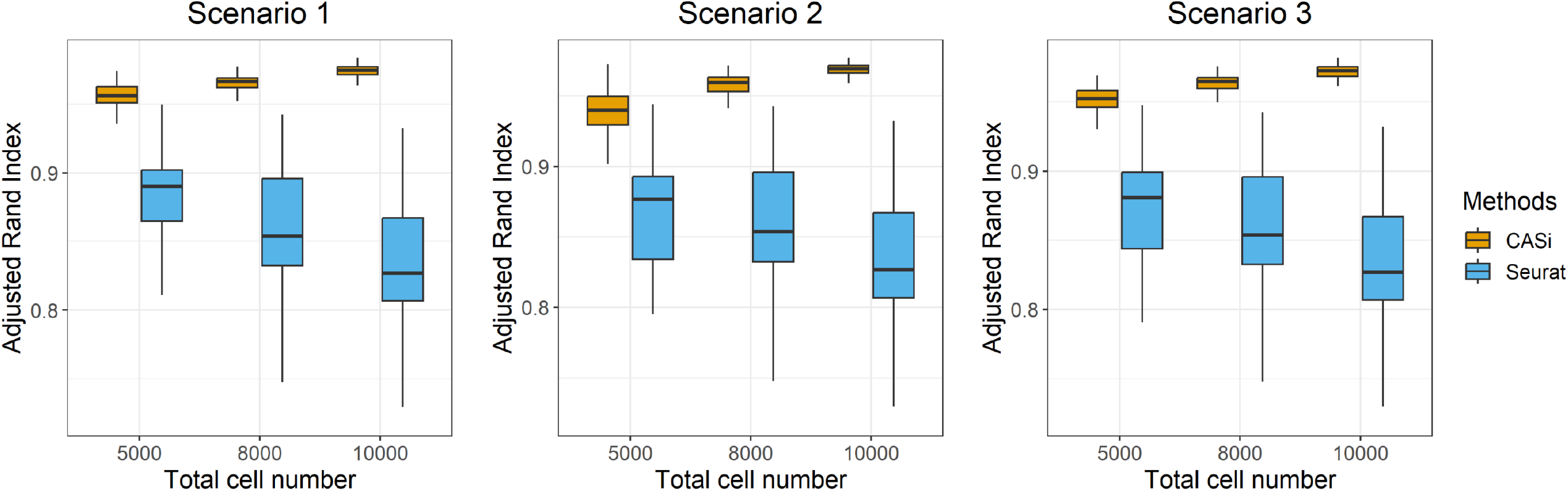
Adjusted rand index (ARI) results of simulation study using different total cell numbers. Based on 200 repetitions, the CASi ARI increases with increasing cell numbers, while the Seurat ARI decreases with increasing cell numbers. This indicates that an overclustering issue does exist in unsupervised clustering methods, such as Seurat, and supervised clustering methods, such as CASi, are able to avoid this issue.

